# Severity-dependent proteomic alterations in the rat hippocampus following pilocarpine-induced status epilepticus

**DOI:** 10.64898/2026.01.08.698453

**Authors:** Surabhi Soni, Yibo Li, Season K. Wyatt-Johnson, Uma K. Aryal, Amy L. Brewster

**Affiliations:** Department of Biological Sciences, Southern Methodist University, Dallas, TX; Department of Psychological Sciences, Purdue University, West Lafayette, IN; Department of Comparative Pathobiology, Purdue University College of Veterinary Medicine, West Lafayette, IN; Biochemistry and Molecular Biology, Oklahoma State University, Stillwater, OK

**Author notes:** **Corresponding Author:** Amy L. Brewster, Ph.D. Associate Professor Department of Biological Sciences Dedman College of Humanities & Sciences Southern Methodist University 6501 Airline Rd, Dallas, TX 75205 Office: DLSB 240; Lab DLSB 215/219. Authors contributed equally (Surabhi Soni, Yibo Li).

## Abstract

Status epilepticus (SE) is a prolonged seizure state that can induce lasting hippocampal damage and promote the development of spontaneous seizures and cognitive deficits. The severity and duration of SE strongly influence these long-term outcomes; however, many experimental studies rely on behavioral assessments such as the Racine scale, which may not capture subclinical or non-convulsive seizure activity. Consequently, the molecular consequences of lower-severity seizures that do not meet conventional criteria for severe SE, but may nonetheless contribute to epileptogenesis, remain poorly understood. To address this gap, we investigated whether behavioral seizure severity in the pilocarpine model correlates with distinct proteomic alterations in the hippocampus. Seizures were induced in adult male rats using pilocarpine, and animals were behaviorally classified into three groups: control, mild SE, and severe SE. Hippocampal tissue was collected from control (n = 3), mild SE (n = 5), and severe SE (n = 6) rats and subjected to mass spectrometry-based proteomic analysis. Proteomic profiles were analyzed using partial least squares discriminant analysis (PLS-DA), and differentially expressed proteins (DEPs) were identified using volcano plots. Functional enrichment analyses were performed using Gene Ontology (GO) and Kyoto Encyclopedia of Genes and Genomes (KEGG) pathway databases. Our findings revealed distinct hippocampal proteomic signatures across control, mild SE, and severe SE groups, as revealed by PLS-DA. Severe SE was associated with widespread proteomic alterations, including 129 DEPs linked to synaptic structure, RNA regulation, and metabolic processes. In contrast, mild SE was associated with fewer changes (81 DEPs), primarily involving synaptic organization and endocytosis. A direct comparison of the severe and mild SE groups identified 76 DEPs enriched in pathways related to synaptic plasticity and neurodegeneration. Notably, 23 proteins showed a stepwise expression pattern across groups, suggesting a molecular gradient correlated with seizure severity. Correlation analyses identified and further confirmed glial and inflammatory molecules as candidate molecular markers associated with seizure burden. In conclusion, behavioral seizure severity in the pilocarpine model corresponds to distinct hippocampal proteomic profiles, with severe SE inducing broader and more pronounced molecular alterations than mild SE. Importantly, lower-severity SE is associated with biologically meaningful changes in pathways related to synaptic organization and cellular processing, highlighting mechanistic differences between mild and severe SE that may contribute to epileptogenic progression.

## Introduction

A prolonged and ongoing seizure lasting more than five minutes can be life-threatening and, if not promptly controlled, may result in long-term neurological consequences. This condition, known as status epilepticus (SE), is defined by the International League Against Epilepsy (ILAE) as “*a condition resulting either from the failure of the mechanisms responsible for seizure termination or from the initiation of mechanisms, which lead to abnormally prolonged seizures [after time point t1 (i.e., 5 mins)]. It is a condition which can have long-term consequences [after time point t2 (i.e., 30 mins)], including neuronal death, neuronal injury, and alteration of neuronal networks, depending on the type and duration* (*1*).*”* In clinical settings, SE episodes that are successfully terminated with anti-seizure medications (ASMs) within 30 minutes are associated with lower morbidity and mortality (2). In contrast, SE that persists beyond 30 minutes increases the risk of developing epilepsy along with additional long-term neurological problems, including cognitive impairment (2).

Evidence from pre-clinical models further supports that, beyond its immediate clinical risks, SE induces widespread molecular, cellular, structural, and physiological changes in susceptible brain areas such as the hippocampus (3–5), which can lead to the generation of hyperexcitable neural circuits that promote spontaneous recurrent seizures (SRS) and impair hippocampal-dependent cognitive functions (6, 7). In rodent models, SE can be induced using chemoconvulsants such as the muscarinic receptor agonist pilocarpine or the kainate receptor agonist kainic acid, and later terminated pharmacologically with ASMs such as diazepam (6). Prolonged SE episodes lasting more than 60 minutes are associated with greater neuronal injury and an increased risk for subsequent development of SRS and cognitive decline (7–10). Because these models recapitulate key aspects of SE pathophysiology and can lead to acquired temporal lobe epilepsy (TLE) similar to that observed in humans (6, 7), chemoconvulsant-induced SE paradigms are widely used in mechanistic studies aimed at identifying causal pathways and therapeutic targets.

Although EEG is the most objective method for determining seizure severity and confirming the onset of SE in experimental models, many studies rely on behavioral scoring systems such as the Racine scale (6, 7, 11). This widely used and accepted scale categorizes seizure severity from mild facial automatisms (Stage 1) to generalized tonic-clonic seizures with loss of posture (Stage 6), which is the most severe stage (11). While the pathological consequences of prolonged, high-severity SE are well documented in our work and others (3–5, 12–16), considerably less is known about the molecular impact of prolonged lower-severity seizures (i.e., continuous level 3) that do not meet the behavioral thresholds typically used to define severe SE (e.g., ≥ Stage 4 for ≥30 minutes) in rodents. This knowledge gap limits our ability to determine whether mild but sustained seizure activity engages pathological mechanisms distinct from, or overlapping with, those triggered by severe SE. To address this, we investigated how behavioral seizure severity, operationally defined as sustained Racine Stage 3 (“mild SE”) versus ≥ Stage 4 (“severe SE”) for one hour, shapes the hippocampal proteome, a brain region highly vulnerable to SE-induced cellular, molecular, and physiological alterations.

## Materials and Methods

### Animals

Sprague Dawley rats (males, 175-200g; Envigo) were kept at room temperature with unrestricted access to food and water (24 hours a day, 7 days a week). The rooms followed a 12-hour light-dark cycle (6:00-18:00). Rats were housed in pairs at the Psychological Sciences animal facility at Purdue University. All procedures adhered to Protocol #1309000927, approved by the Purdue Animal Care and Use Committee, and complied with NIH and institutional guidelines (17).

### Pilocarpine-induced SE

Rats were pretreated with scopolamine methyl bromide (1 mg/kg; Cat# S8502-1G, Sigma-Aldrich) 30 minutes before receiving pilocarpine (280 mg/kg; i.p.; Cat# P6503-10G, Sigma-Aldrich) following previously described protocols (3, 12, 14). Control rats received 0.9% saline (i.p.). Seizures were scored according to a modified Racine scale as follows: 1 = rigid posture, mouth movements; 2 = tail clonus; 3 = partial body clonus with head bobbing; 4 = rearing with severe whole-body clonic seizures while retaining posture; 5 = rearing and falling; and 6 = tonic-clonic seizures with jumping or loss of posture. Rats that progressed to stage 3 and maintained these seizures for up to 1 hour were classified as the *mild SE* group. Rats that reached stage 4 or higher and sustained those behavioral seizure manifestations for 1 hour were classified as the *severe SE* group. At 1 hour from seizure onset, SE was terminated with diazepam (10 mg/kg, i.p.; Hospira, Inc.). Following SE induction, all rats received 0.9% saline (i.p.) for hydration and were given supplemental nutrition (chocolate Ensure and Kellogg’s Fruit Loops) for up to three days post-SE. Body weight was measured daily over 10 days. Final rat counts: control, n = 3; mild SE, n = 5; severe SE, n = 6.

### Tissue preparation

Two weeks after pilocarpine-induced seizures, rats were euthanized with a lethal dose of Beuthanasia (200 mg/kg, i.p.) and transcardially perfused with ice-cold 1X phosphate-buffered saline (PBS; 137 mM NaCl, 2.7 mM KCl, 4.3 mM Na₂HPO₄, 1.47 mM KH₂PO₄, pH 7.4). Hippocampi were immediately dissected, frozen, and stored at −80 °C for proteomic analysis. The two-week time point after pilocarpine was selected based on the robust hippocampal neuronal, synaptodendritic, inflammatory, and glial changes previously observed following SE in our previous studies (3–5, 16).

### Proteomics sample preparation and data analysis

Hippocampal tissues were processed at the Purdue Proteomics Core Facility at the Bindley Bioscience Center, as previously described (18). Briefly, tissues were suspended in 100 mM ABC and homogenized in Precellys homogenization vials (Bertin Technologies SAS, France) for 90 seconds at 6500 rpm. Protein concentration was measured with the bicinchoninic acid assay (Pierce Chemical Co., USA). Fifty μg of protein were precipitated at −20 °C with acetone overnight. Then, samples were centrifuged at 13,500 × g for 15 minutes at 4 °C. The pellets were briefly dried in a vacuum centrifuge without heat on, resuspended in 10 µL of 10 mM DTT in 8 M urea, and incubated at 37 °C for 1 hour. An equal volume of alkylating mixture (195 mL acetonitrile, 1 mL triethyl phosphate (TEP), 4 mL Iodoethanol) was added. Samples were incubated at 37 °C for an additional hour, dried in a vacuum centrifuge, and digested with Lys-C/Trypsin in 50 mM ammonium bicarbonate (ABC) at a 1:25 enzyme-to-protein ratio using a barocycler (60 cycles: 20 kPSI for 50 seconds and 1 ATM for 10 seconds, at 50 °C). Digested peptides were desalted using C_18_ Silica MicroSpin Columns (The Nest Group, Inc. USA). Eluted and cleaned peptides were dried in SpeedVac and resuspended in 3% acetonitrile/96.9% MilliQ, and 0.1% formic acid (FA) at a final concentration of 1mg/ml.

### Liquid Chromatography-Tandem Mass Spectrometry (LC-MS/MS) analysis

Peptides were analyzed in a Dionex UltiMate 3000 RSLC nano System (Thermo Fisher Scientific, Odense, Denmark) coupled with an Orbitrap Fusion Lumos Tribrid Mass Spectrometer (Thermo Fisher Scientific, Waltham, MA, USA) (19). Reverse phase peptide separation was accomplished using a trap column (300 mm ID ’ 5 mm) packed with 5 mm 100 Å PepMap C18 medium coupled to a 50-cm long × 75 µm inner diameter analytical column packed with 2 µm 100 Å PepMap C18 silica (Thermo Fisher Scientific) at 50°C. Mobile phase solvent A: 2% acetonitrile (ACN), 98% water, and 0.1% Formic Acid (FA). Mobile phase solvent B: 80% ACN, 20% water, and 0.1% FA. One mg of peptide sample was loaded to the trap column in a loading buffer (3% acetonitrile, 0.1% FA) at a flow rate of 5 mL/min for 5 min and eluted from the analytical column at a flow rate of 200 nL/min using a 160-min LC gradient as: 6.5 to 27% of solvent B in 82 min, 27-40% of B in next 8 min, 40-100% of B in 7 min at which point the gradient was held at 100% of B for 7 min before reverting to 2% of B, and hold at 2% of B for next 15 min for column equilibration. The column was washed and equilibrated by using three 30-minute LC gradients before injecting the following samples. All data were acquired on an Orbitrap mass analyzer and collected using an HCD fragmentation scheme. For MS scans, the scan range was 350-1600 m/z at a resolution of 120,000. The automatic gain control (AGC) target was set to 4 × 10^5^, the maximum injection time was 50 ms, dynamic exclusion lasted 30 seconds, and the intensity threshold was 5.0 × ^104^. MS data were acquired in Data Dependent mode with a cycle time of 5s/scan. MS/MS data were collected at a resolution of 15,000.

### Data Analysis

LC-MS/MS data were analyzed using MaxQuant software (version 1.6.3.3) against the combined non-redundant protein sequence database from Uniprot (www.uniprot.org) for rat, for protein identification and label-free relative quantitation. The following parameters were used for database searches: precursor mass tolerance of 10 ppm; enzyme specificity of trypsin/Lys-C, allowing up to 2 missed cleavages; oxidation of methionine (M) as a variable modification, and iodoethanol (C) as a fixed modification. False discovery rate (FDR) of peptide spectral match (PSM) and protein identification was set to 0.01. Proteins with LFQ # 0 and MS/MS (spectral counts) ≥ 2 were considered true identifications and used for downstream statistical analysis and data visualization.

### Bioinformatics analysis

For statistical analyses, Label-Free Quantification (LFQ) intensities were employed, and the dataset was limited to proteins quantified in at least half of the samples in each condition. The missing values were imputed by using K-nearest neighbors (KNN), and normalization was performed. Volcano plot, Partial Least Squares Discriminant Analysis (PLS-DA), and hierarchical cluster analysis were performed using R software (version 4.4.0). For hierarchical clustering, Euclidean distance was used as the distance measure, and complete linkage was used to combine clusters. The ShinyGO 0.76, a web-based tool (http://bioinformatics.sdstate.edu/go/), was used to perform enrichment analysis of differentially expressed proteins (DEPs), enabling the study of gene ontology (GO) and Kyoto Encyclopedia of Genes and Genomes (KEGG) pathways. The GO classification consisted of biological processes (BP), molecular functions (MF), and cellular components (CC). *P-value* < 0.05 was set as the threshold of significance. For KEGG pathway enrichment analysis, DEP information was mapped to the KEGG database to identify enriched pathways. Spearman correlation coefficients (ρ) and corresponding p-values were calculated in R (version 4.4.0) using the cor.test() function, with a two-tailed significance threshold. Protein abundance values were log2-transformed and median-normalized across samples before correlation analysis to minimize technical variation and improve comparability across animals.

### Statistical analysis

The differences between the two groups were evaluated using the Student’s *t*-test, uncorrected. FDR correction (Benjamini-Hochberg) was performed and was included in the supplementary files. The plots were generated in R (version 4.4.0) and GraphPad Prism software (version 10.2.0). Differences were considered statistically significant at *p* < 0.05. Significantly altered proteins are included in Supplementary Table 1.

## Results

The Racine scale is a standardized scoring system used to classify the severity of behavioral seizures in rodent models of epilepsy, especially during and after chemically induced SE. It was first developed in the 1970s by Ronald Racine and remains widely used in epilepsy research to evaluate seizure progression in rodent models of SE (6, 7, 11). The Racine scale includes the following stages: Stage 1 involves rigid posture and mouth movements; Stage 2 includes tail clonus; Stage 3 features partial body clonus and head nodding; Stage 4 is characterized by rearing (standing on hind limbs) with forelimb clonus; Stage 5 involves rearing and falling; and Stage 6 refers to tonic-clonic seizures with jumping or loss of posture (Fig. 1B). This scale helps researchers estimate seizure severity and progression after the chemoconvulsant administration. Although various modifications and expanded versions have been developed to incorporate additional behaviors or seizure types, the original Racine scale remains a key tool for assessing seizure severity in studies using rats and mice. Across our studies, about 90% of pilocarpine-treated rats develop and maintain seizures above Racine scale 4, which we classify as severe SE (Fig. 1C). These animals are included in subsequent analyses (3–5, 12, 14, 15).

**Figure 1.**
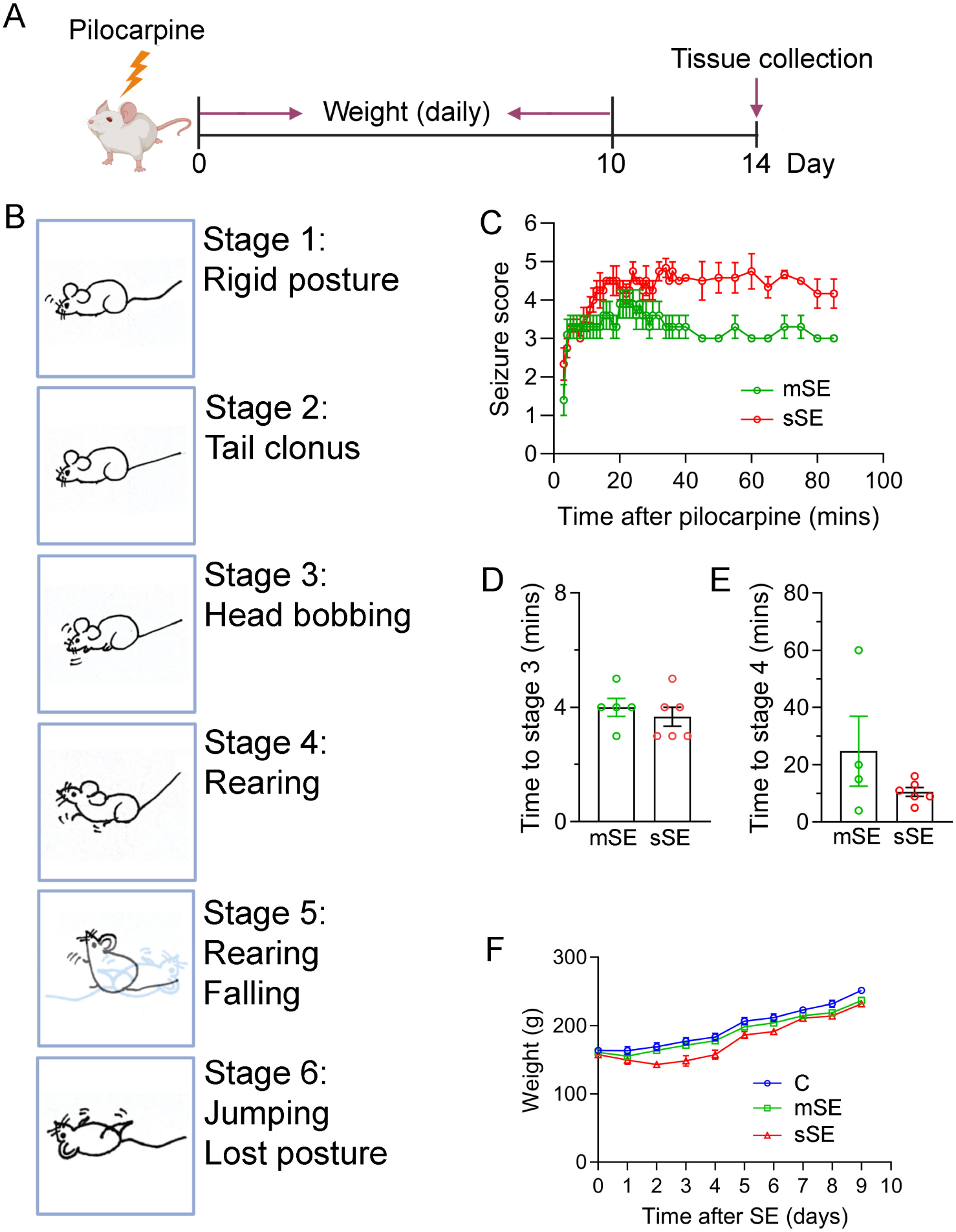
Characterization of seizure induction. (A) Experimental timeline. Pilocarpine was administered on Day 0 to induce seizures. Body weight was monitored daily for 10 days, and tissues were collected on Day 14. (B) Schematic representation of Racine seizure scale stages. Stage 1: Rigid posture, mouth moving; Stage 2: Tail clonus; Stage 3: Partial body clonus, head nodding; Stage 4: Rearing (standing on hind limbs) with forelimb clonus; Stage 5: Rearing and falling; Stage 6: Tonic-clonic seizure with jumping or loss of posture. (C) Seizure scores over time following pilocarpine injection in the mild SE and severe SE groups are shown. (D-E) Quantification of latency to reach Racine Stage 3 (D) and Stage 4 (E) in mild SE and severe SE groups. (F) Body weight trajectories during the 10 days following pilocarpine injections. Abbreviations: mild SE, mSE; severe SE, sSE.

However, during our studies, we also noticed that a small group of animals (∼5%) given the same pilocarpine dose initially reached a Racine scale 3-4 but did not progress to higher levels. Instead, their behavioral seizures remained at Racine scale 3 consistently for at least one hour (Fig. 1C), which we classified as mild SE. When we compared daily body weight among the mild SE, severe SE, and control groups, we found that rats in the severe SE group showed significantly reduced body weight between days 2 and 6 after pilocarpine treatment (p < 0.05). In contrast, rats in the mild SE and control groups did not lose weight and exhibited comparable weight gain throughout the monitoring period (Fig. 1F).

To determine whether the observed differences in behavioral seizure severity are reflected in hippocampal alterations, we analyzed the hippocampal proteome at 2 weeks after SE (Fig. 2A), when we had previously found robust and significant changes in dendritic and glial markers (3–5, 12, 13, 16). Partial least squares discriminant analysis (PLS-DA) of the proteomic data revealed clear separation among the three groups (Fig. 2B), and there were significant differences in both component 1 (Fig. 2C) and component 2 (Fig. 2D) across all groups, suggesting distinct proteomic profiles associated with SE severity as measured by the Racine scale.

**Figure 2.**
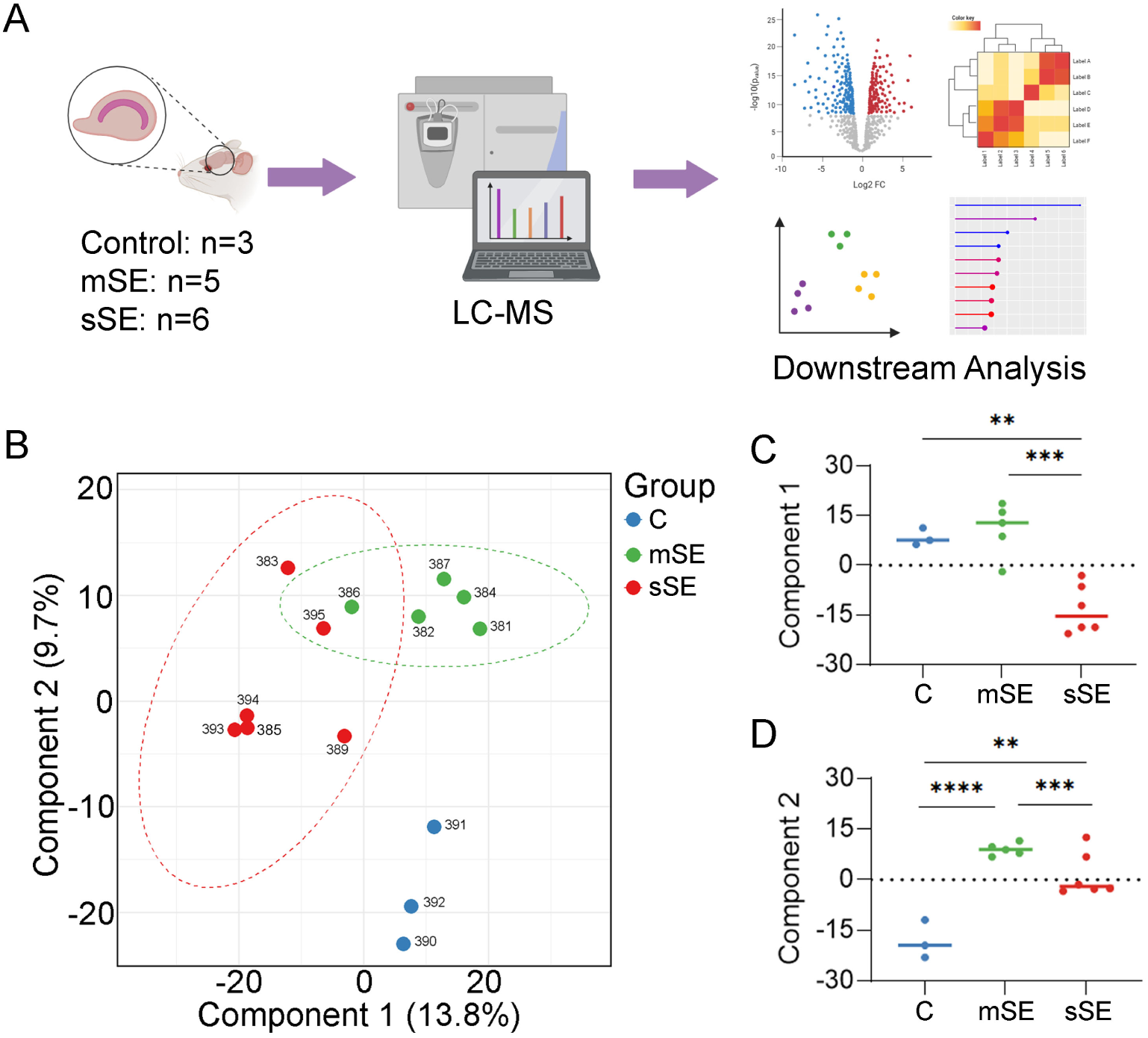
Diagram of the experimental workflow. (A) Rat hippocampal tissues were collected from each group (control, *n* = 3; mild SE, *n* = 5; severe SE, *n* = 6). Proteins were quantified using liquid chromatography-tandem mass spectrometry (LC-MS), followed by statistical analysis, functional enrichment, and pathway analysis to identify differentially expressed proteins and enriched pathways. (B) PLS-DA plot showing separation of Control, mild SE, and severe SE groups based on proteomic expression data. Each point represents an individual sample, and group clustering indicates distinct molecular signatures associated with seizure severity. Ellipses represent 95% confidence intervals for each group. (C-D) Component 1 and Component 2 scores from the PLS-DA showing significant differences among control, mild SE, and severe SE groups. Abbreviations: Control, C; mild SE, mSE; severe SE, sSE.

A direct comparison between the control and severe SE groups revealed distinct global proteomic profiles (Fig. 3A). A total of 129 DEPs were identified, with 72 up-regulated and 57 down-regulated in the severe SE group relative to the control group (Fig. 3B). The heatmap of DEPs comparing the severe SE and control groups, showed a hierarchical group-specific clustering (Fig. 3C). GO enrichment analysis of the DEPs revealed significant changes in the abundance of proteins associated with cytoskeletal protein binding, cortical cytoskeleton, synapses, and regulation of mRNA metabolic processes (Fig. 3D). KEGG pathway enrichment analysis indicated the involvement of pathways such as PI3K-Akt signaling, phagosome formation, and metabolic processes including cysteine and methionine metabolism (Fig. 3E). These results suggest that severe SE induces widespread long-lasting alterations in the hippocampal proteome with notable changes involving the regulation of RNA and synaptic structures as well as in metabolic and intracellular signaling pathways.

**Figure 3.**
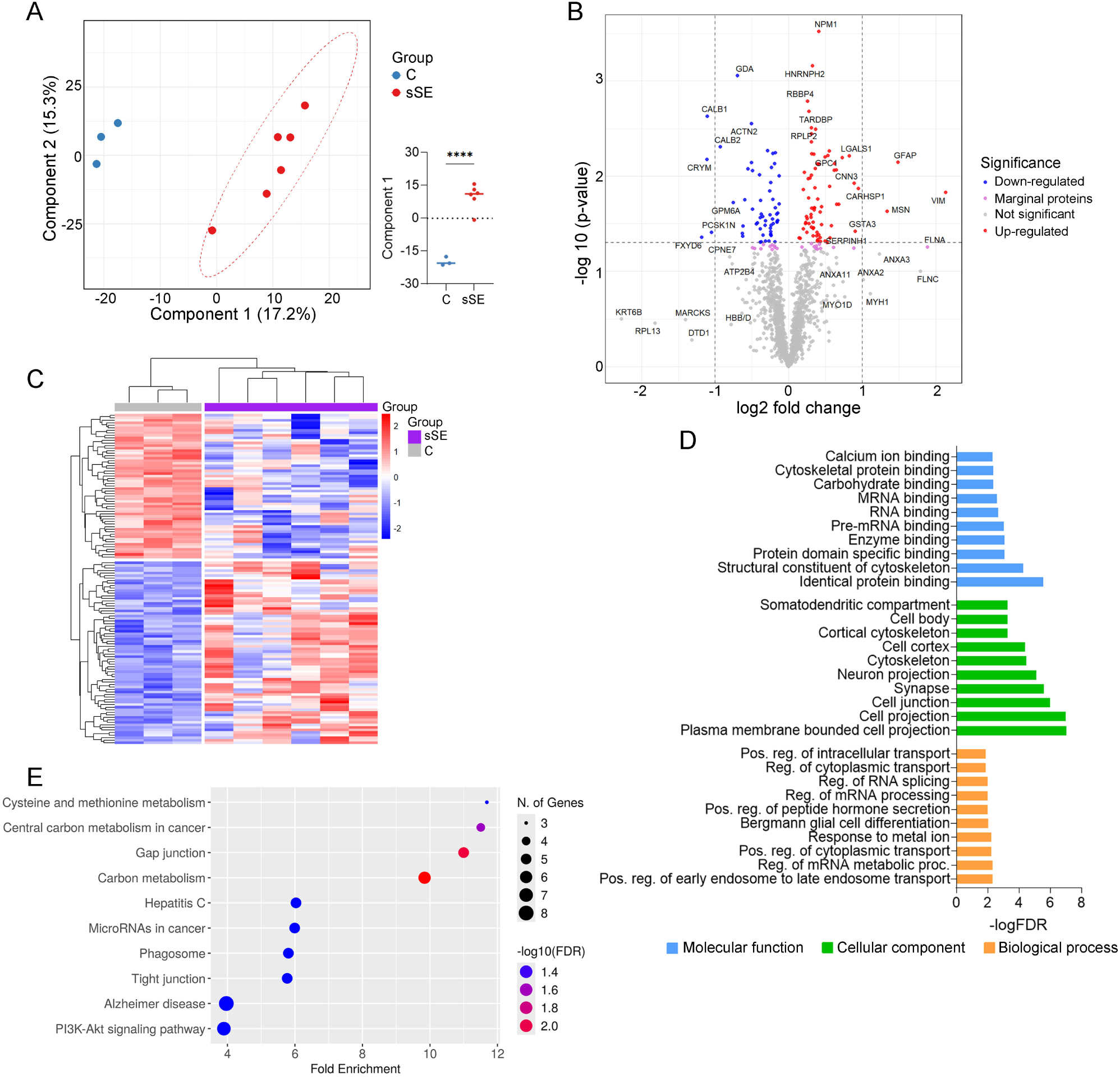
Proteomic differences between the Control and severe SE groups. (A) PLS-DA plot shows clear separation between Control and severe SE groups based on global protein expression profiles. Each point represents an individual sample. Ellipses denote 95% confidence intervals. (B) Volcano plot displaying differentially expressed proteins between the Control and severe SE groups. Red and blue dots indicate significantly up- and downregulated proteins, respectively (p < 0.05). (C) Heatmap of differentially expressed proteins clustered by expression pattern across samples. Rows represent proteins; columns represent individual samples. (D) Gene Ontology (GO) enrichment analysis of DEPs, categorized by Biological Process (BP), Cellular Component (CC), and Molecular Function (MF), the top 10 are shown. (E) KEGG pathway enrichment analysis of DEPs showing the top 10 significantly enriched pathways. Abbreviations: Control, C; severe SE, sSE.

Next, we compared the control and mild SE groups. PLS-DA analysis revealed a clear separation between the two groups (Fig. 4A), suggesting distinct hippocampal proteomic profiles. A total of 81 DEPs were identified, with 63 up-regulated and 18 down-regulated in the mild SE group (Fig. 4B). The heatmap of these DEPs also showed distinct expression patterns between these groups (Fig. 4C). GO enrichment analysis showed that most of the altered proteins were associated with biological processes related to synaptic organization, protein localization, and cytoskeleton regulation, and molecular functions involved in cytoskeletal, receptor, and nucleotide binding functions (Fig. 4D). KEGG pathway analysis identified endocytosis as the only significantly enriched pathway (Fig. 4E).

**Figure 4.**
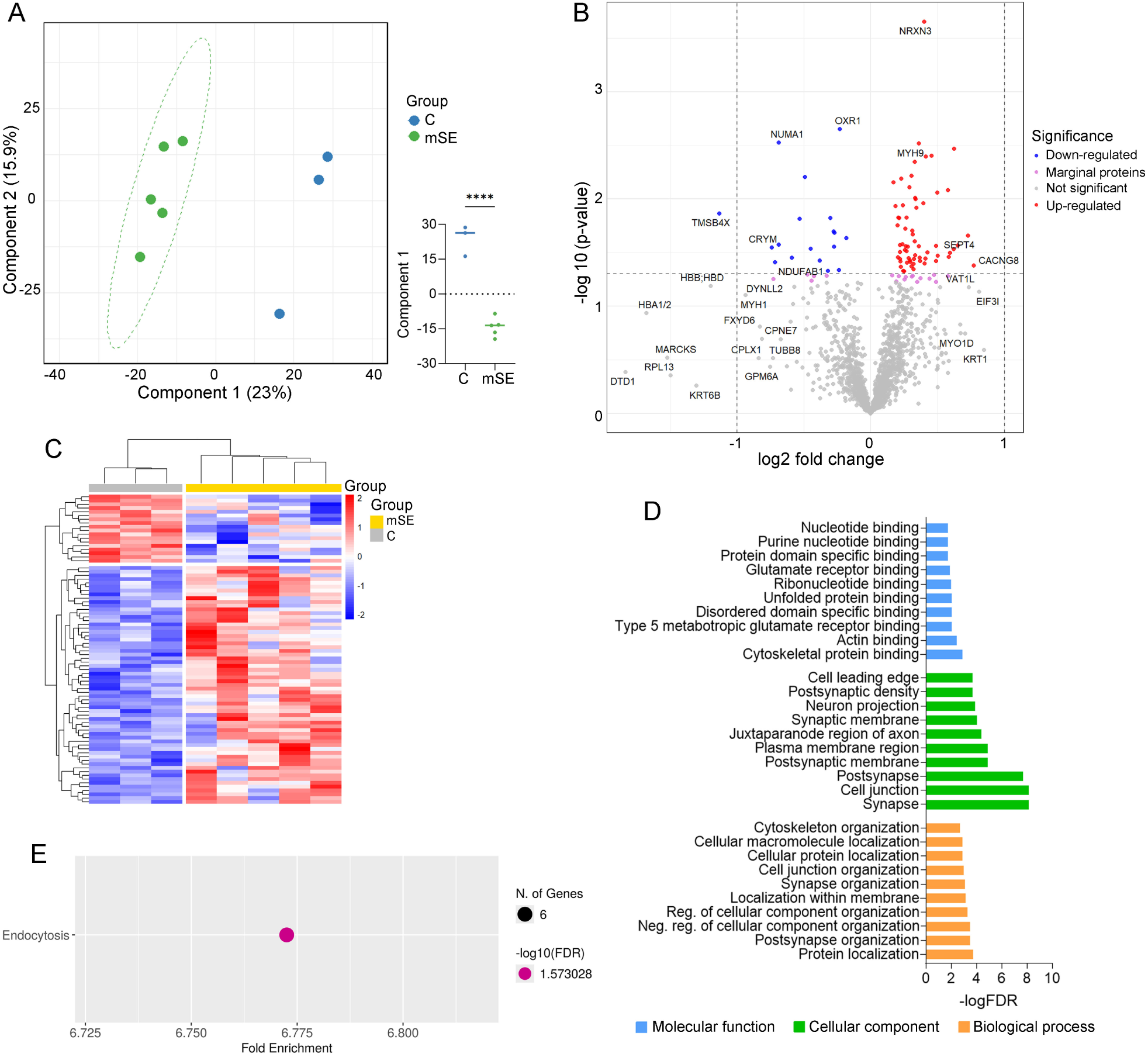
Proteomic differences between the Control and mild SE groups. (A) PLS-DA plot showing clear separation between Control and mild SE groups based on global protein expression profiles. Each point represents an individual sample. Ellipses denote 95% confidence intervals. (B) Volcano plot displaying differentially expressed proteins between the Control and mild SE groups. Red and blue dots indicate significantly up- and downregulated proteins, respectively (p < 0.05). (C) Heatmap of differentially expressed proteins clustered by expression pattern across samples. Rows represent proteins; columns represent individual samples. (D) Gene Ontology (GO) enrichment analysis of DEPs, categorized by Biological Process (BP), Cellular Component (CC), and Molecular Function (MF), the top 10 are shown. (E) KEGG pathway enrichment analysis of DEPs. Abbreviations: Control, C; mild SE, mSE.

To study proteomic changes related to SE severity, we compared the mild SE group with the severe SE group. (Fig. 5). PLS-DA analysis showed a clear and significant separation between the groups (Fig. 5A). Statistical analysis (*p* < 0.05) identified 76 DEPs, with 37 up-regulated and 39 down-regulated in the severe SE group (Fig. 5B), as evidenced by distinct clustering and expression patterns in the heatmaps (Fig. 5C). These differences may reflect a molecular transition that may contribute to epileptogenesis in this model. GO enrichment analysis revealed that DEPs are associated with synaptic function and plasticity (BP), enriched in synaptic structures (CC), as well as with actin, receptor, calcium, and adhesion-related binding activities (MF) (Fig. 5D). KEGG pathway analysis revealed enrichment in synaptic and neurodegenerative pathways, including mitophagy, ubiquitin-mediated proteolysis, cell adhesion, and diseases such as Alzheimer’s and Parkinson’s diseases, and amyotrophic lateral sclerosis (Fig. 5E). These findings suggest that progressing to, and sustaining, a severe SE episode at Racine scale level 4 or above leads to widespread synaptic remodeling and activation of neurodegenerative pathways, compared with sustained Racine scale level 3 seizures.

**Figure 5.**
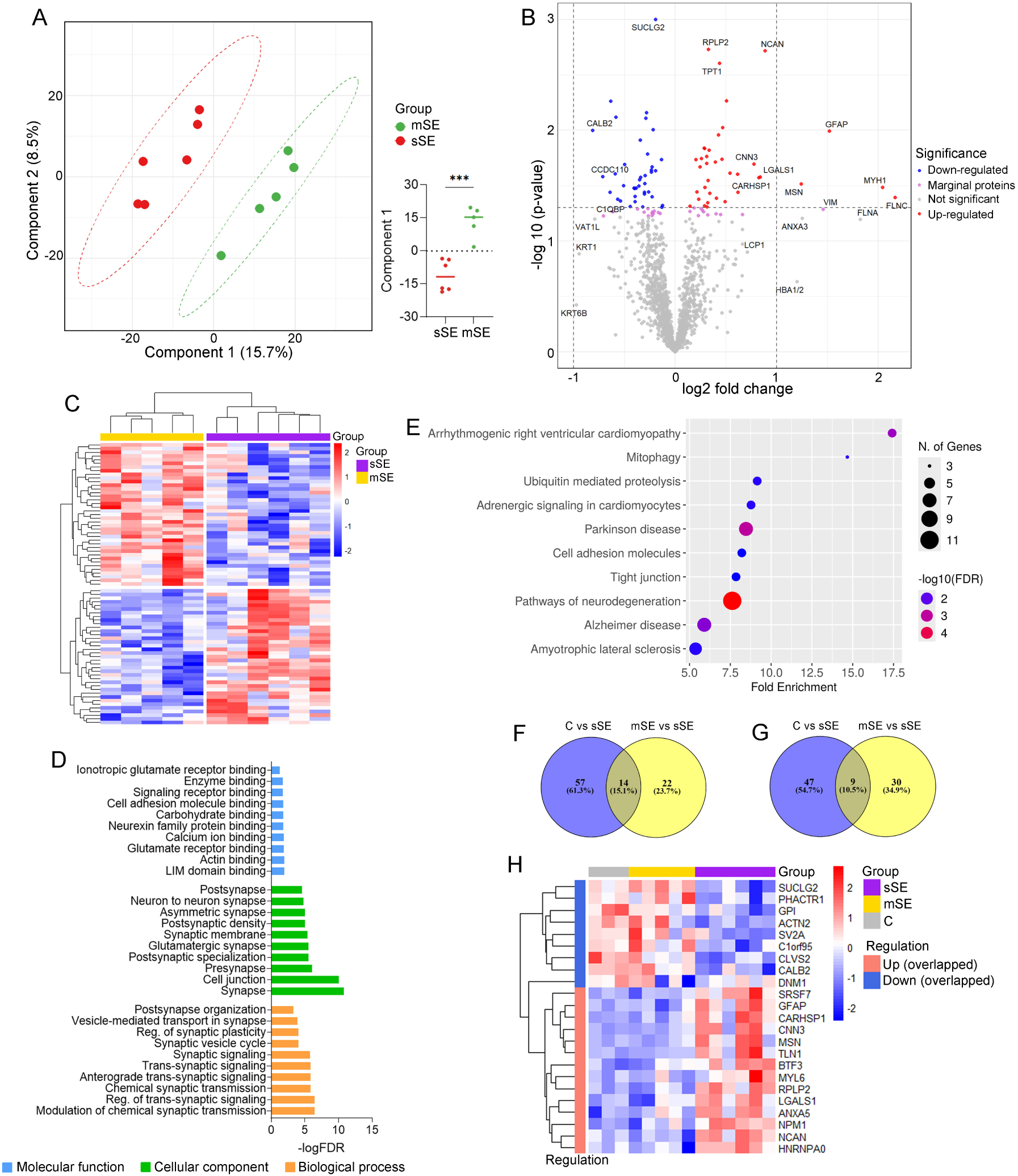
Proteomic differences between the mild SE and severe SE groups. (A) PLS-DA plot showing clear separation between mild SE and severe SE groups based on global protein expression profiles. Each point represents an individual sample. Ellipses denote 95% confidence intervals. (B) Volcano plot displaying differentially expressed proteins between severe SE and mild SE groups. Red and blue dots indicate significantly up- and downregulated proteins, respectively (p < 0.05). (C) Heatmap of differentially expressed proteins clustered by expression pattern across samples. Rows represent proteins; columns represent individual samples. (D) Gene Ontology (GO) enrichment analysis of DEPs, categorized by Biological Process (BP), Cellular Component (CC), and Molecular Function (MF), the top 10 are shown. (E) KEGG pathway enrichment analysis of DEPs showing the top 10 significantly enriched pathways. (F) Venn diagrams showing 14 up-regulated proteins were commonly altered in both the Control vs severe SE and mild SE vs severe SE comparisons. (G) Venn diagrams showing 9 down-regulated proteins commonly altered in both Control vs severe SE and mild SE vs severe SE comparisons. (H) Heatmap of the 23 overlapped proteins (14 up-regulated and 9 down-regulated) identified in both comparison analyses. Rows represent proteins and columns represent individual samples from the Control, mild SE, and severe SE groups. Protein expression values are Z-score normalized. Abbreviations: mild SE, mSE; severe SE, sSE.

To identify proteins consistently associated with SE severity and potential epileptogenesis, we compared DEPs between control vs. severe SE and mild SE vs. severe SE groups. We identified 14 proteins that were significantly upregulated (Fig. 5F) and 9 that were significantly downregulated (Fig. 5G) exclusively in the severe SE group. Proteins with increased abundance in severe SE included GFAP (Glial Fibrillary Acidic Protein), LGALS1 (Galectin-1), NCAN (Neurocan), and ANXA5 (Annexin A5), which provide structural support in glial cells and can promote astrocyte activation and immune-matrix signaling (LGALS1), remodel the extracellular matrix and regulate synaptic stability and plasticity (NCAN), and reflect membrane stress, calcium dysregulation, and apoptosis-associated signaling (ANXA5).

Collectively, increased levels of these proteins suggest ongoing SE-induced reactive astrogliosis, extracellular matrix reorganization, and neuroinflammation. These processes are known to enhance neuronal hyperexcitability and contribute to seizure propagation and persistence in epilepsy (20–22). In addition, increases in MSN (Moesin), CNN3 (Calponin 3), TLN1 (Talin 1), and MYL6 (Myosin Light Chain 6), which support actin-based adhesion, contraction, and motility, indicate cytoskeletal reorganization and mechanotransductive signaling (23). Together, changes in these proteins in the hippocampus suggest that reactive astrocytes or glial cells undergo pronounced structural remodeling in response to severe SE. Conversely, downregulated proteins specific to severe SE included DNM1 (Dynamin 1), SV2A (Synaptic Vesicle Glycoprotein 2A), PHACTR1 (Phosphatase and Actin Regulator 1), ACTN2 (Alpha-Actinin 2), and CLVS2 (Clavesin 2), all of which are linked to synaptic structure and function, may contribute to the extensive synaptodendritic loss observed in the hippocampus two weeks post-SE in this model (4, 5, 12, 15).

To further explore the relationship between protein abundance and SE severity, we performed Spearman correlations between 33 top proteins (C vs. severe SE, p < 0.05, |log2FC| > 0.5) and four seizure metrics: maximum Racine score, cumulative seizure duration, and times to stages 3 and 4. The heatmap (Fig. 6A) highlights these relationships. GFAP, VIM (Vimentin), and NCAN showed strong positive correlations with seizure severity, linking higher levels to more severe SE. GFAP, VIM, and NCAN correlated strongly with both seizure duration (Fig. 6B-D) and Racine score (Fig. 6E-G). In contrast, SRR (Serine Racemase), ACTN2, and CALB2 (calretinin), negatively correlated with seizure parameters (Fig. 6H-J), suggesting that reduced hippocampal levels of proteins involved in synaptic organization, cytoskeletal stability, and calcium buffering may be associated with altered neuronal homeostasis and increased network excitability that may be pro-epileptogenic. Taken together, these proteomic findings suggest that proteins linked to gliosis, neuroinflammation, structural remodeling, and neuroprotection are associated with seizure severity in this SE rat model.

**Figure 6.**
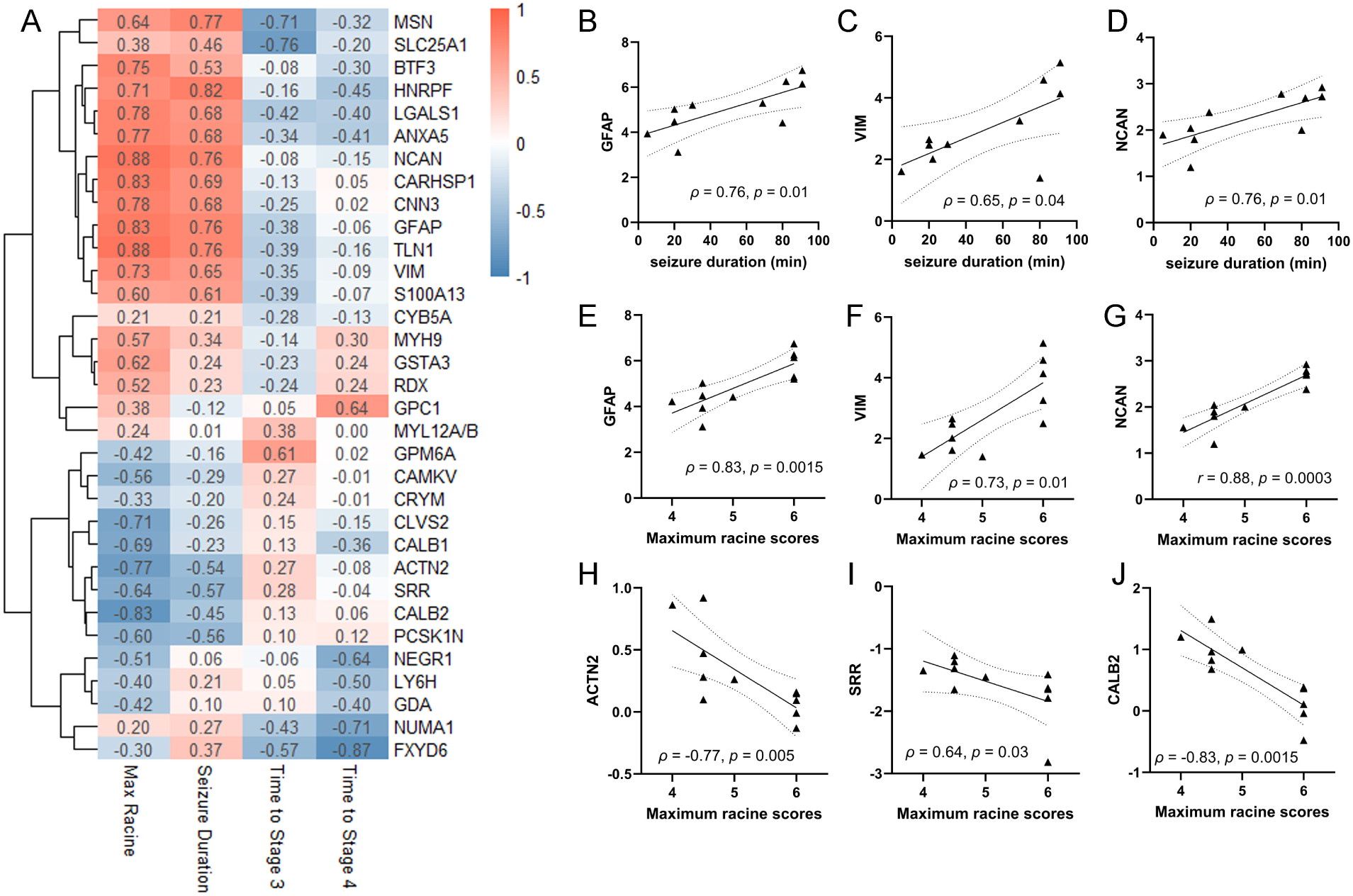
Correlations between the top 33 DEPs and seizure parameters. (A) Heatmap showing Spearman correlation coefficients (ρ values) between the normalized abundance of the top 33 proteins (C vs severe SE, p < 0.05, |log2FC| > 0.5) and four seizure metrics: maximum Racine score, cumulative duration, and times to stages 3 and 4. Red indicates positive correlations; blue indicates negative correlations. (B-J) Scatter plots show the relationship between seizure severity parameters and the abundance of the proteins GFAP, VIM, NCAN, ACTN2, SRR, and CALB2. Each panel represents one protein.

## Discussion

This study employed a well-characterized pilocarpine-induced SE rat model to define hippocampal proteomic signatures across three behavioral outcomes: no seizures (Control), mild SE, and severe SE. Prior work has largely compared severe, sustained SE to controls, leaving the molecular impact of shorter, non-progressing seizures (Racine ≤3) poorly defined. Inclusion of the mild SE group enabled detection of intermediate proteomic changes and revealed a graded molecular response aligned with seizure severity. PLS-DA demonstrated clear biological separation among all three groups, indicating that behavioral classification reflects distinct hippocampal proteomic states. Severe SE produced the most extensive alterations, with 129 DEPs enriched in pathways related to synaptic architecture, RNA regulation, and metabolism.

Mild SE was associated with 81 DEPs involving synaptic organization and endocytosis, representing a distinct molecular profile rather than a reduced version of the severe SE response. Direct comparison of severe versus mild SE identified 76 DEPs enriched in synaptic plasticity and neurodegeneration pathways, suggesting these proteins may mark early mechanistic transitions in epileptogenesis.

Our comparison of SE and control groups aligns with previous proteomic reports in SE models, which show distinct alterations between the groups (24–26). A study by Bitsika et al. conducted a hippocampal proteomic analysis 30 days after intrahippocampal kainate-induced SE in male mice and identified 175 differentially expressed proteins (97 upregulated and 78 downregulated) (24), which match the expression patterns observed in the pilocarpine model. After GO analysis, this study reported that proteins related to neuron projection (e.g. Purkinje cell protein 4, Tenascin), synaptic plasticity and synaptic organization (e.g., Map1b/a,

Paralemmin-1, MAPK) were consistently downregulated, similar to our observations following pilocarpine-induced SE. The study also highlighted pronounced alterations in astrocyte- and glia-related proteins accompanying the development of chronic epilepsy (24). Further supporting the involvement of glial and neuronal remodeling, Li et al. performed a proteomic analysis of the hippocampal dentate gyrus in male mice four weeks after pilocarpine-induced SE (25). They identified SE-driven changes in proteins associated with metabolic, synaptic, structural, and transcriptional regulatory pathways. Specifically, within the dentate gyrus, metabolism-related (e.g., L-lactate dehydrogenase B, pyridoxal phosphatase) and transcription-related proteins (e.g., nascent polypeptide-associated complex alpha subunit isoform b, heterogeneous nuclear ribonucleoprotein D-like) were downregulated, whereas structural (e.g., profilin-1, vimentin) and synaptic proteins (e.g., synuclein) were upregulated. Together, these results further emphasize the coordinated metabolic, structural, and glial changes accompanying the progression to chronic epilepsy.

Interesting differences were noted in synaptic and metabolic pathway changes between the above-mentioned proteomic studies using whole hippocampal homogenates (24) and those using dissected dentate gyrus (DG)(25). The loss of synaptic elements in whole hippocampal homogenates may reflect the pronounced loss of synaptodendritic structures within the CA1-CA3 regions, which has been widely reported after SE across models of acquired epilepsies (27). In contrast, analysis of the DG alone showed increases in synaptic-related pathways (25), which may be due to enhanced connectivity among granule cells and mossy cells, and to mossy fiber sprouting associated with epilepsy progression in rodent models (28–32). These findings support the idea that using the entire hippocampus in omics studies can dilute region-specific effects. Thus, while the proteomic findings support broad changes across multiple pathways, identifying which changes may cause post-SE epilepsy and which are secondary to the SE-induced injury requires further region-specific cell-level analysis.

By incorporating a mild SE cohort, this study extends prior work by capturing a frequently overlooked intermediate seizure phenotype. Proteomic analysis of this group showed enrichment in synaptic organization and endocytosis. These shifts suggest that mild seizures can disturb synaptic trafficking and cytoskeletal control, but they may not be sufficient on their own to trigger subsequent epilepsy. We also identified 23 proteins uniquely associated with the severe SE phenotype. In particular, GFAP, VIM, and NCAN demonstrated strong positive correlations with pilocarpine-induced seizure duration and higher Racine scale scores. This finding aligns with previous studies, which have well established GFAP and VIM as markers of astrocyte activation and reactive gliosis, hallmarks of seizure-induced neuroinflammation and damage in both SE and chronic epilepsy models (20, 33). NCAN, an extracellular matrix (ECM) protein, has also been implicated in seizure-related ECM remodeling, a process increasingly recognized as contributing to epileptogenesis (34–36). While we did not directly quantify neuronal loss or gliosis histologically in the present study, the proteomic signatures observed are consistent with well-characterized structural, glial, and inflammatory responses to severe SE in this same model (3–5, 12, 15).

The finding of strong negative correlations between seizure severity and the expression of ACTN2, SRR, and CALB2 suggests that these proteins may serve as molecular markers of seizure-induced neuronal dysfunction. ACTN2, a cytoskeletal protein involved in synaptic structure, was significantly downregulated with increasing seizure severity. This aligns with previous findings of reduced α-actinin-2 levels during dendritic remodeling in the DG gyrus after SE (37), supporting its role in synaptic destabilization during epileptogenesis. SRR, which synthesizes D-serine, also showed a strong negative correlation with seizure severity. This is consistent with reports that exogenous D-serine mitigates neuronal loss and inflammation in epilepsy models (38), suggesting that reduced SRR may impair endogenous neuroprotection. CALB2, a calcium-binding protein, was similarly decreased with greater seizure severity, in line with evidence of selective calretinin-positive neuron loss following kainic acid-induced SE (39).

Together, these findings highlight ACTN2, SRR, and CALB2 as potential markers of seizure burden, reflecting synaptic disruption and vulnerability that may be epileptogenic. However, it is important to note that unprovoked behavioral seizures were not monitored, and EEG analysis was not performed in the rats used in this study. Therefore, it is possible that the rats assigned to the severe SE group may have had subsequent electrographic ictal or interictal activity that contributed to the more pronounced proteomic changes observed at 2 weeks post-SE. Similarly, it remains unclear whether rats in the mild SE group developed epilepsy. To fully capture the potential of mild versus severe SE to generate epilepsy and associated molecular changes in the hippocampus, continuous EEG monitoring would be required.

Another limitation of this study is that, after multiple-comparison adjustment, few proteins remained statistically significant; therefore, our analysis was conducted using uncorrected values. This limitation is common in discovery-based mass spectrometry proteomic studies, particularly those involving complex tissues and small sample sizes, where stringent FDR correction can be overly conservative, thereby overlooking biologically relevant changes. In our prior work using a comparable proteomic workflow, proteins that did not survive FDR correction were subsequently validated using independent approaches, such as western blotting, supporting the biological relevance of these findings (18). Nevertheless, as discussed above, our findings are comparable to other proteomics studies in similar rodent models of SE.

An additional consideration from this study is the need for clearly defined, standardized criteria to classify experimental groups based on seizure severity after pilocarpine-induced SE. Relying only on behavioral signs using the Racine scale alone might be inadequate, as behavioral severity does not always mirror the actual electrographic seizure burden. Accordingly, experimental protocols should explicitly specify which Racine scale stages are used to distinguish mild from severe SE for downstream analyses. EEG studies support that SE severity is associated with differences in electrographic activity patterns and hippocampal molecular profiles in rodent models of SE and acquired TLE (40–44). Therefore, analyzing animals that reach sustained mild versus severe SE as separate groups, when classification is based only on behavioral criteria, may reduce within-group variability and facilitate interpretation of experimental outcomes. In contrast, pooling animals across SE severities is likely to increase variability and obscure severity-dependent effects.

In summary, this study demonstrates that seizure severity in an initial pilocarpine-induced SE episode is reflected in distinct hippocampal proteomic signatures, with mild and severe SE exhibiting potentially graded synaptic, metabolic, and glial alterations that may underlie differential trajectories of epileptogenesis. Although these results offer biologically significant insights that could guide new mechanistic targets, they are currently restricted to males. Given previous findings of sex-specific proteomic and transcriptomic differences in both a genetic mouse model of epilepsy and human epilepsy (18, 45, 46) these results may not fully apply to females, highlighting the need for future studies to examine both sexes.

## Author Contributions

SS analyzed data, created figures, and wrote the manuscript. YL analyzed data, created figures, and wrote the manuscript. SJ performed animal experiments and collected tissues. UA conducted LC-MS/MS. ALB designed the study and wrote the manuscript.

## Acknowledgments

The author(s) acknowledge the use of the facilities of the Bindley Bioscience Center, a core facility of the NIH-funded Indiana Clinical and Translational Sciences Institute. This research was supported by NS096234 (ALB); Hamilton Undergraduate Research Scholars Program (ALB & SS), Southern Methodist University.

## Data availability

Raw data from this study are available from the Texas Data Repository (Accession No. UBOM3D).

## Supplementary tables

Supplementary Table 1: Significantly altered proteins in each comparison.

Supplementary Table 2: Correlation between 33 selected top proteins and four seizure metrics.

